# Overactive PDGFRα and PDGFRβ promote distinct yet overlapping phenotypes of skeletal muscle fibrosis and stiffness, with PDGFRβ also driving drastic muscle growth

**DOI:** 10.64898/2026.01.02.697406

**Authors:** Mariola Gimla, Szczepan Olszewski, Jacob L. Brown, Christiana Raymond-Pope, Sandra Rigsby, Frederick F. Peelor, Hae Ryong Kwon, Longbiao Yao, Lorin E. Olson, Jordan D. Fuqua, Benjamin F. Miller

## Abstract

**Background:** Fibrosis accumulates in skeletal muscle over time and leads to greater muscle rigidity, stiffness, and increased risk of injuries. However, investigations of experimental models to study the mechanisms through which muscle fibrosis occurs are often confounded by injury or disease. The contribution of platelet-derived growth factor receptors alpha and beta (PDGFRα or PDGFRβ) to muscle fibrosis is yet to be clarified. We hypothesized that both receptors would promote ECM deposition and fibrosis, causing muscle stiffening and weakness, with sex-specific differences arising due to hormonal influences on receptors. **Methods:** To test this hypothesis, we used a mouse model with inducible overactive PDGFRα or PDGFRβ signaling and assessed various indicators of muscle function, metabolism, motor coordination, exercise capacity, collagen deposition, and muscle stiffness. **Results:** Overactive PDGFRα led to more collagen deposition, increased collagen crosslinking, and higher AGE/LOX protein levels, all of which correlated with greater muscle stiffness compared to CON. Overactive PDGFRβ resulted in greater muscle mass and lower fat mass, and had higher collagen deposition in female mice compared to CON. There were also sex-specific differences with fibrotic remodeling, muscle stiffness, and muscle size in response to overactive PDGFRα and PDGFRβ signaling. **Conclusion:** These findings establish PDGFRα and PDGFRβ signaling as distinct regulators of muscle remodeling and establish overactive PDGFRα as a mouse model to study skeletal muscle fibrosis in the absence of other confounding variables.

## INTRODUCTION

Skeletal muscle extracellular matrix (ECM) is essential for muscle structure as it provides a framework for myofibers, nerve supply, capillaries, and other supporting cells (1). Muscle ECM also plays an essential role in force transmission (2, 3) and mechanotransduction (4–6). Although ECM consists of multiple components such as glycoproteins, proteoglycans, and glycosaminoglycans, the predominant protein component is collagen. With age and with certain metabolic disorders (i.e., insulin resistance) there is increased collagen abundance, increased stiffness, and increased collagen crosslinking (7). These changes result in muscle fibrosis leading to stiffer and less functional skeletal muscle.

There are three types of cells that produce and secrete collagen in mouse skeletal muscle: fibroblasts, fibro-adipogenic progenitors (FAPs), and skeletal muscle progenitor cells (8). Platelet derived growth factor receptors (PDGFR) alpha (α) and/or beta (β) are found on each of these three cells and can promote their deposition of collagen in different tissues (9–15). During development, elevated PDGFRα signaling in mice leads to increased connective tissue growth in multiple organs, including skeletal muscle, leading to excessive and widespread fibrosis (16–18). Although PDGFRβ is less studied than PDGFRα, PDGFRβ lineage cells have been shown to shift to a fibrotic phenotype in skeletal muscles of aged mice (10). Despite the progress in understanding PDGFR signaling, the relative contributions of overactive PDGFRα or PDGFRβ activity in promoting excessive deposition of collagen and fibrosis during aging is unknown.

A major limitation in advancing our understanding, and therefore treatment, of age-related muscle fibrosis is the availability of appropriate experimental models to study the mechanisms through which muscle fibrosis occurs. Animal models used to study muscle fibrosis often include dystrophic mouse models such as the *mdx* mouse, a model of Duchenne Muscular Dystrophy (1, 19), or non-dystrophic injury models that include lacerations, tenotomy, crushing, or chemically-induced injury (1). Each of these models presents other confounding variables (e.g., severe muscle wasting or injury-induced trauma) that limit applicability to mechanisms responsible for age-related fibrosis. Therefore, animal models are needed that facilitate the independent effects of fibrosis on muscle without affecting other aspects of muscle health.

The goal of this study was to determine whether overactive PDGFRα or PDGFRβ leads to skeletal muscle fibrosis and impaired function. We hypothesized that both overactive PDGFRα and PDGFRβ would result in skeletal muscle fibrosis, muscle stiffness, and impaired muscle function. Because estrogen has been shown to induce PDGFR expression (20), we also hypothesized that there would be sex-related differences in these two models. Secondarily, we sought to identify a new model to study the mechanisms of muscle fibrosis that was not complicated by muscle damage or a disease model. To study these hypotheses and goals, we used two different inducible mouse models that express a gain-of-function of either PDGFRα or PDGFRβ. While there were overlapping outcomes between the two mutant models, we found that overactive PDGFRα leads to significant muscle fibrosis and stiffness while overactive PDGFRβ was not fibrotic and had a robust increase in muscle mass and a decrease in fat mass. Each model also showed sex-related differences in multiple physiological outcomes like collagen crosslinking, treadmill exercise capacity, force production, and muscle stiffness. Whereas the overactive PDGFRa mouse could be useful for studying fibrosis independent of muscle damage or disease, the overactive PDGFRb has a hypertrophic phenotype that may be useful for other mechanistic studies.

## METHODS

### Animal models

We maintained all mouse strains on a mixed C57BL/129 genetic background. The bKFlp and aRFlp alleles were made in 129 genetic background ES cells, while the remaining alleles have a C57BL6 background. All animal comparisons were age-matched, and we used littermate controls whenever possible. We used both male and female mice to test the effect of sex. To create the tamoxifen-inducible PDGFRα gain-of-function model, we bred *Ubc-CreER^tg^* (JAX:007001) (21) and *Pdgfra^K-Flp^* mice (JAX:038627) (22). To create the tamoxifen-inducible PDGFRβ gain-of-function model, we bred mice carrying *Pdgfra^CreER^*(JAX:032770) (23) and *Stat1^flox^* (JAX:032054) (24) alleles with *Pdgfrb^K-Flp^* mice. *Stat1^flox^* was included in the PDGFRβ gain-of-function model because previous work has shown that elevated PDGFRβ signaling leads to early lethality due to excessive Stat1-induced inflammation (25). The *Pdgfrb^K-Flp^* knock-in/knock-out mice have a floxed stop cassette and cDNA encoding a constitutively active *Pdgfrb* gene (D849V; PDGFRβ^K^), with a T2A-*flpo* sequence replacing the PDGFRβ^K^ stop codon. We prepared tamoxifen (Tmx) as a 20mg/ml stock in corn oil. To induce Cre recombination at 4-months-old, we gavaged mice three times with 100 mg Tmx/kg bw. We housed mice in the vivarium at the Oklahoma Medical Research Foundation under the care of the Department of Comparative Medicine. We group-housed animals with *ad libitum* access to food and water in a room on a 12:12 h light:dark cycle under constant temperature and humidity control. The Institutional Animal Care and Use Committee (IACUC) of the Oklahoma Medical Research Foundation approved all procedures performed on mice. For ease of reading, PDGFR mutant mice will be identified as PDGFR+ and control mice will be identified as CON.

### Study design

After induction of recombination at 4 months of age, the mice were aged to 7 months when they underwent a battery of tests that was spread out over 2 months to limit animal stress (**Figure 1A**). Most mice were used for all outcomes, however, some muscle preps were compromised during *ex vivo* assays, so we do not have data for all mice for those assays. Specific sample sizes are indicated in figure legends. Seven days before the tissue collection, we injected mice intraperitoneally (i.p.) with a bolus of 99% D_2_O, equivalent to 5% of the body water pool (7, 26, 27). Following i.p. injection animals had *ad lib* access to 8% D_2_O-enriched drinking water. Prior to harvest, we measured *in vivo* muscle contractile function. After euthanasia with CO_2_ and cervical dislocation we collected blood through cardiac puncture and harvested and weighed lower limb skeletal muscles. We then put the right extensor digitorum longus muscle (EDL) in oxygenated Ringer’s solution and measured *ex vivo* muscle stimulation and passive stiffness. All other right limb muscles were trimmed of fat and connective tissues, and flash-frozen in liquid nitrogen. Left limb muscles were prepared for histology. We also measured tibia length and collected bone marrow and plasma. Physiological measurements, histology, microscopy, and morphological analyses were performed with the investigator blinded.

**Figure 1.**
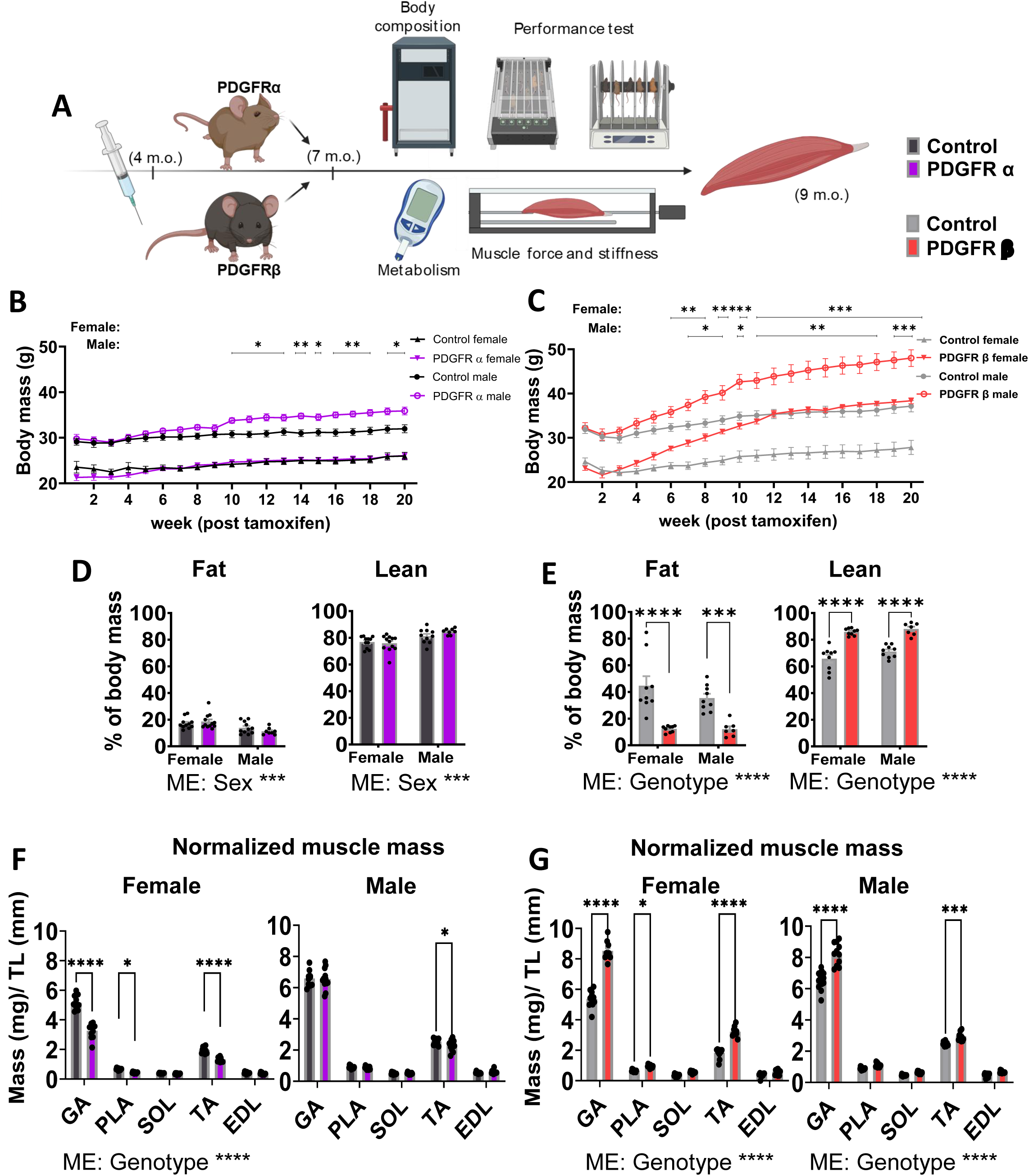
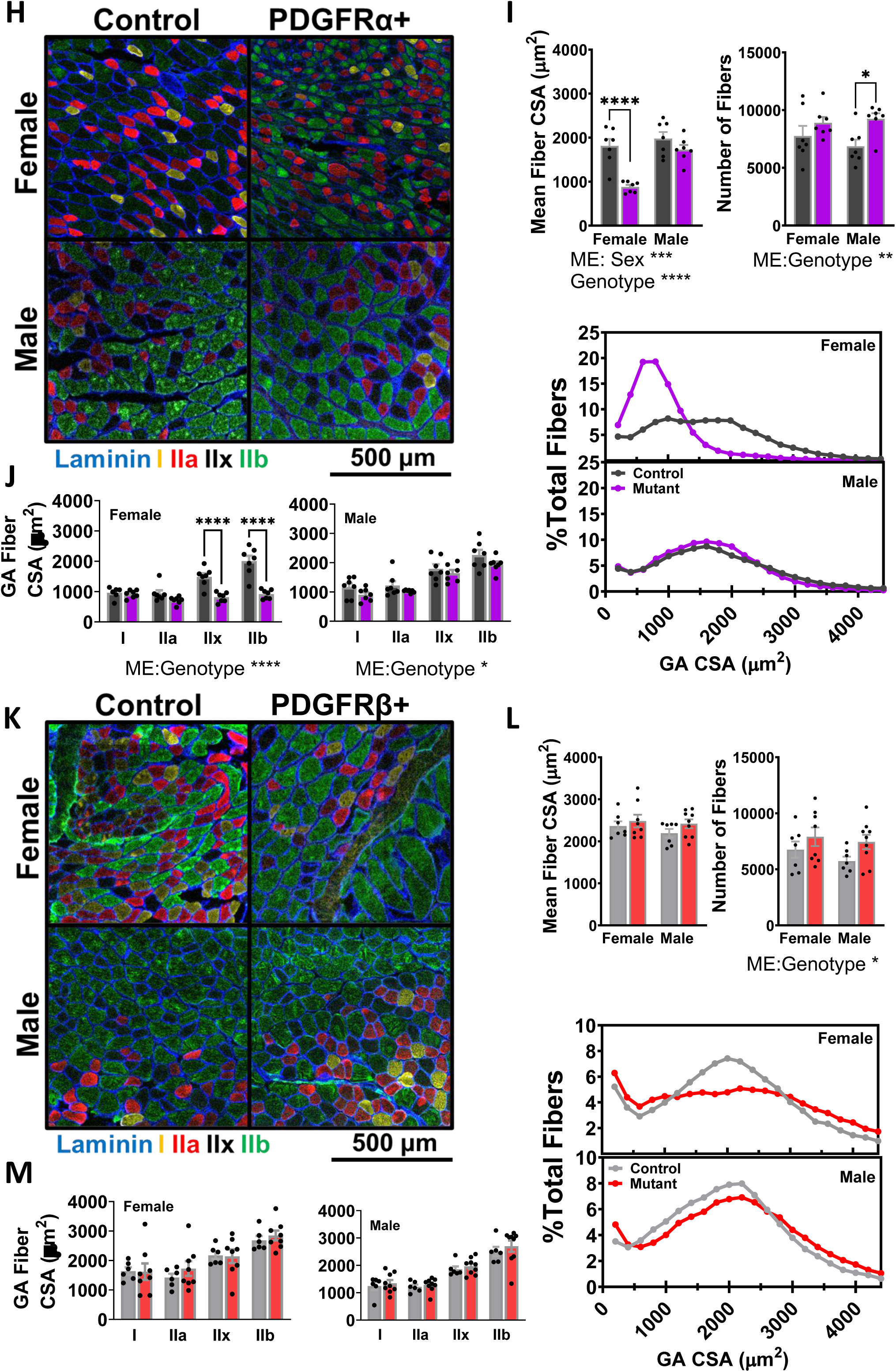
Mice with overactive PDGFRβ signaling have greater body mass, lean mass, and less fat mass. **(A)** Study design for PDGFRα and PDGFRβ mice. Changes in body mass over length of intervention for **(B)** PDGFRα and **(C)** PDGFRβ mice. Percentage of fat and lean mass in **(D)** PDGFRα and **(E)** PDGFRβ mice (*n=7-12*). Individual wet muscle mass (GA, PLA, SOL, TA, and EDL) normalized to tibia length from **(F)** PDGFRα and **(G)** PDGFRβ mice (*n=7-9*). Representative images of MHC fiber type staining, myofiber mean cross-sectional area (CSA), total fiber number, CSA per fiber type, and distribution of fiber size in **(H, I, J)** PDGFRα and **(K, L, M)** PDGFRβ mice (*n=7-8*). Gastrocnemius (GA), plantaris (PLA), soleus (SOL), tibialis anterior (TA), and extensor digitorum longus (EDL). Data are means ± SEM. *p<0.05, **p<0.01, ***p<0.001, ****p<0.0001.

### Body composition

We measured body composition at 7 months of age using EchoMRI™-500 Body Composition Analyzer (Echo Medical Systems, Houston, TX, USA). Mice were placed in a ventilated plastic canister while being held immobile during assessments.

### Endurance capacity

We acclimated mice to the treadmill (Treadmill Simplex II, Columbus Instruments, Columbus, OH, USA) for three days by running at 10 m/min for 10 minutes. On the fourth day, animals were tested with an initial speed of 10 m/min for 10 minutes at a 5% incline. At this point, we gradually increased the speed by 2 m/min until the mice are exhausted. We calculated the total work and max power performed during the treadmill run:

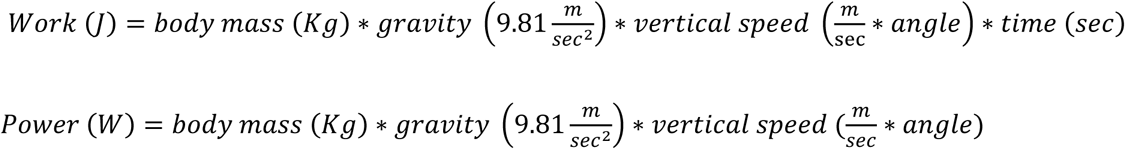

We calculated max power by using the max vertical velocity which occurred at the time just prior to failure. An electric grid and brush located at the end of the treadmill were used to encourage the animals to run. We terminated the test when a mouse stopped responding to tail brushing/electric grid continuously for 3 seconds. We assessed blood lactate and blood glucose before and at the termination of the test by tail bleeding.

### Rotarod

We placed mice in home cages in the testing room for at least 1 hour before testing to minimize the effects of stress on behavior during testing. We acclimated mice to the rotarod (Rota-Rod ENV-577 | ENV-577M, Med Associates, Fairfax, VT, USA) for three days. On the first day, mice were placed on the apparatus at a speed of 4 rpm for 60 seconds in 6 different trials with 5 minutes rest between trials. Next day for 4 rpm x 4 sets and 10 rpm x 4 sets for 60 seconds each. On the last day of acclimatization, 4 rpm x 4 sets and 20 rpm x 4 sets for 60 seconds each. We tested animals on the fourth day with an initial speed of 4 rpm accelerated to 40 rpm in 5 minutes. The test was terminated when the animal falls off the rotating rod. We repeated the procedure for a total of four trials separated by 5 minutes intervals.

### *In vivo* muscle contractile function

We assessed muscle function in plantar flexors by stimulating the tibial nerve, according to our previously published methods (28). In summary, the knee of the mouse was secured between pivots, and the foot was placed in and secured to the foot plate attached to a force transducer. Two needle electrodes (monopolar PTFE coated stainless steel) (Chalgren Enterprises, Gilroy, CA) were used for percutaneous stimulation of tibial nerve. Tetanic isometric contractions at optimal length were elicited by stimulations of 150 Hz for 300 msec using the 1300A: 3-in-1 Whole Animal System (Aurora Scientific Inc, Aurora, ON, Canada). We collected and analyzed data to determine maximal muscle torque using the Dynamic Muscle Analysis software (ASI 611A v.5.321, Aurora Scientific Inc, Ontario, Canada). Additional contractile parameters from the tetanic curve were also evaluated, including average rates of contraction and relaxation and the time to maximal contraction.

### *Ex vivo* skeletal muscle specific force

We conducted studies of limb muscle isometric function as previously described (29, 30). Mice were sacrificed, and the EDL muscle was excised and immediately placed in oxygenated Krebs buffer for dissection. One end of the muscle was tied to a Dual-Mode Muscle Lever System (300C-LR, Aurora Scientific Inc, Aurora, Canada) and the other end onto a hook. Muscles were placed at optimal length and allowed 10 min of thermoequilibration to the desired temperature (32 °C). Thereafter, measurements of maximal force were initiated. In all electrical stimulations, a supramaximal current (600–800 mA) of 0.25 ms pulse duration was delivered through a stimulator (701 C, Aurora Scientific Inc.), while train duration for isometric contractions was 300 ms. We collected and analyzed all data using commercial software (DMC and DMA, Aurora Scientific).

### D_2_O labeling and protein synthesis

We measured protein synthesis by determining deuterium incorporation into protein of gastrocnemius muscles as previously described (7, 27, 31–33). To measure the synthesis of subcellular fractions, we homogenized and then fractionated skeletal muscle tissue according to our previously published procedures (7, 27, 31–33). We then ran derivatized samples of alanine and ran them on an Agilent 7890A GC coupled to an Agilent 5975C MS. We determined body water enrichment on serum using a Liquid Water Isotope Analyzer (LWIA-45-EP, Los Gatos Research, San Jose, CA). We used rates of tracer incorporation to calculate protein fractional synthesis rates (FSR, %/day) as performed previously in our lab (7, 27, 31–33).

#### Glucose regulation

We performed glucose tolerance tests (GTT) after 6h of fasting, ∼2pm (fasting started ∼8am). Briefly, we transferred animals to clean cages without food at the beginning of the fasting period, and the test was initiated with a baseline assessment of glycaemia via tail bleeding with glucometer (CONTOUR^®^NEXT EZ, Ascensia Diabetes, New Jersey USA). Subsequently, we performed an i.p. injection of a glucose solution prepared in saline (2 g/kg body weight). We measured blood glucose at 15, 30, 60, and 120 min after the glucose injection. We performed insulin tolerance tests (ITT) after 4h of fasting, ∼2 pm (fasting started ∼10am). We assessed baseline levels of blood glucose and then i.p. injected insulin solution prepared in saline (0.75 U/kg of body weight; Humulin R, Eli Lilly, Indianapolis, IN). We measured blood glucose at 15, 30, and 60 min after insulin injection. On a separate day, we transferred mice to clean cages without food and then measured blood glucose with a glucometer (CONTOUR^®^NEXT EZ, Ascensia Diabetes, New Jersey USA) at the end of a 16-hour fasting period at 20:00.

### Pepsin collagen solubility and hydroxyproline assay

To quantify the proportions of mature cross-linked collagen and non-cross-linked and immature cross-linked collagen in gastrocnemius muscles, we treated ∼30 mg of tissue with pepsin as previously described (7). We determined the collagen concentration in the pepsin-soluble and pepsin-insoluble fractions using a hydroxyproline (OHP) assay as we previously described (7). We measured concentration through absorbance reading at 558 nm with the BioTek Synergy H1 microplate reader (Agilent Technologies Santa Clara, CA, US).

### Quantification of pyridinium cross-links

To quantify pyridinium crosslinks, we measured pyridinoline and deoxypyridinoline (P&D) using the MicroVue PYD EIA enzyme immunoassay (Quidel Corporation: San Diego, CA 92121, USA) following the manufacturer’s instructions. We measured concentration through absorbance reading at 405 nm with the BioTek Synergy H1 microplate reader (Agilent Technologies Santa Clara, CA, US).

### Quantification of myostatin

To measure myostatin in plasma we used a GDF11-K01 mouse myostatin/GDF8 ELISA Kit (Eagle Biosciences, Inc., Nashua, NH 03063, USA) following the manufacturer’s instructions. We measured concentration through absorbance reading at 450 nm with the BioTek Synergy H1 microplate reader (Agilent Technologies Santa Clara, CA, US).

### Immunoblot analysis

Skeletal muscle samples were prepared, processed, and analyzed as described previously (27). We prepared primary antibody dilutions as follows: lysyl oxidase (LOX) (Novus Biologicals, NBP2-24877, 1:1000), advanced glycation end products (AGEs) (Abcam, ab23722, 1:500), total p70S6 kinase (Cell Signaling Technology [CST] 9202, 1:1000) and phospho-p70S6K (Thr389) (CST 9205, 1:1000), total ribosomal protein S6 (rpS6) (CST 2217, 1:1000) and phospho-rpS6 (Ser235/236) (CST 4858, 1:1000), total 4EBP1 (CST 9452, 1:1000) and phospho-4EBP1 (Thr37/46) (CST 9459, 1:1000). We used primary antibodies with the LI-COR secondary antibody IRDye 800CW Goat anti-Rabbit IgG 926-32211 at 1:10,000 concentration. We normalized results for protein expression to Ponceau stain.

### Histology, fluorescence/confocal microscopy, and myofiber diameter

Upon harvesting, gastrocnemius muscles were directly embedded in tissue frozen medium (Tissue-Tek^®^ O.C.T., Fisher-Scientific) and frozen in liquid nitrogen cooled isopentane for subsequent analysis. We obtained serial sections of muscle samples (10 μm) from the middle section of Gastrocnemius at a temperature of −24–25°C using a Cryotome Leica 3050 (Leica Biosystems, Deer Park, IL, US.)

We determined fiber type and fiber size by post-fixing sections in pre-cooled acetone. Next, we permeabilized muscle sections in PBS with 1% Tween 20, blocked with M.O.M. solution (MKB 2213, Vector Laboratories, Burlingame, CA), rinsed and blocked again in normal goat serum (NGS, Sigma, G-9023) diluted to 5% in PBST (5% NGS), and then incubated overnight with primary antibodies: MHC Type-I (BA-F8, 1:250), Type-IIa (SC-71s, 1:250), Type-IIb (BF-F3s, 1:250) from Developmental Studies Hybridoma Bank (University of Iowa, Iowa City), and Laminin (L9393, 1:500) from Sigma-Aldrich (St. Louis, MO) at 4°C. On the following day, we incubated samples for 1 hour with corresponding isotype specific secondary antibodies in 5% NGS, and then covered with ProLong Diamond Antifade Mountant (Thermo Fisher Scientific). Samples were washed 3 times with PBST after each of the steps outlined above except after the second blocking where no wash was performed. Also, MHC Type-IIx fibers were denoted as “black” fibers (i.e., absence of color/dye). We imaged slides using a Zeiss LSM710 Confocal Microscope (Oberkochen, Germany). We measured muscle fiber cross-sectional area (CSA) using Myovision analysis software (Myovision 2.0 software, University of Kentucky).

To evaluate capillarity, we stained muscle sections with CD31 (Thermo Fisher Scientific, 14-0311-82, 1:100), laminin (Sigma-Aldrich, L9393, 1:500), and DAPI (Invitrogen, 1:10000), using previously described procedures (34). The following corresponding isotype specific conjugated secondary antibodies were used: Alexa Fluor 555 (Invitrogen, A21428, 1:200) and Alexa Fluor 647 (Invitrogen, A21247, 1:200). We then imaged whole tissue sections using a Zeiss Axioscan 7 slide scanner and a 20X objective (Plan-Apochromat 20x /0.8 M27). Images were exported and analyzed using Myovision analysis software (Myovision 2.0 software, University of Kentucky).

To evaluate collagen content and organization (i.e., loosely and densely packed collagen) we stained muscle sections with picrosirius red (PSR) (Abcam, ab246832), as previously described (35). Whole tissue sections were imaged using a Zeiss Axioscan 7 slide scanner with a diascopic cross-linear polarized filter using a 20X objective (EC Plan-Neofluar 20x/0.50 Pol M27). After adjusting white balance and focus parameters, a reference brightfield image and subsequent polarized image of the whole muscle section were captured. We exported images and used Fiji (ImageJ) to conduct thresholding analysis of green and red/yellow collagen staining within the polarized images of each gastrocnemius section. Images were binarized and the area fraction of green (loosely packed) and red/yellow (densely packed) were quantified using the Color Pixel Counter plugin and presented as a percentage of total area.

### Statistical analysis

Results are presented as mean ± *SEM* with individual data points shown for each mouse. Within both PDGFRa and PDGFb experiments we used a two-way analysis of variance (ANOVA) (genotype by sex) followed by Sidaks post hoc analyses where appropriate. Main effects are shown when significantly different. For body mass over time we used a two-way repeated measures ANOVA, followed by Tukey post hoc analyses. Due to limited lanes within a gel, we ran all Western blots with one sex per gel to be able to directly compare between mutant and control animals. We analyzed Western blots with unpaired t-tests. For our correlation between muscle stiffness and collagen we used Pearson R correlation. Values of p<0.05 were considered statistically significant. All statistical tests were performed with GraphPad Prism, v. 10 (GraphPad Software Inc., San Diego, CA).

## RESULTS

We first determined the effects of overactive PDGFRα and PDGFRβ signaling on body composition. We showed that while there was no difference in body mass between female CON and PDGFRα+, body mass was significantly higher in male PDGFRα+ and male and female PDGFRβ+ mice compared to their respective CON (**Figure 1 B and C**) (**Supplemental Figure 1A and B**). We measured body composition with EchoMRI prior to euthanasia and found that there were no differences in lean or fat masses between PDGFRα+ and CON (**Figure 1D**). However, male and female PDGFRβ+ had less % fat mass and greater % lean mass compared to their CON (**Figure 1E**). There was a main effect of genotype for hindlimb muscle mass in the female PDGFRα mice with many of the hindlimb muscles weighing less in PDGFRα+ compared to their CON (**Figure 1F**) (**Supplemental Figure 1 C**). There was also a main effect of genotype for both male and female PDGFRβ mice with many of the hindlimb muscles weighing more in PDGFRβ+ mice compared to CON (**Figure 1G**) (**Supplemental Figure 1D**). Because PDGFR signaling regulates bone growth, we measured tibia length and showed that tibia length was longer in PDGFRβ+ compared to CON while EDL length was also longer in PDGFRα+ in both sexes and in female PDGFRβ+ relative to their respective CON (**Supplemental Figure 1E-H**). Because of this longer bone length, we normalized muscle mass by tibia length (**Figure 1F and G**). For fiber type CSA there was a main effect of genotype for both male and female groups. PDGFRα+ female mutants had lower gastrocnemius (GA) mean myofiber CSA compared to their CON, and this lower CSA was only seen in the fast twitch (i.e., IIx and IIb) muscle fibers (**Figure 1H-J**).There was a main effect of genotype for the total number of fibers in GA muscles with a higher number of fibers in male PDGFRα+ compared to CON (**Figure 1I**). For PDGFRβ mice, both male and female mutants showed a rightward shift in myofiber CSA distribution compared to their respective CON (**Figure 1K-M**). Although there was no significant difference in myofiber CSA there is a main effect of genotype for total number of fibers in GA muscles, and this combined with the rightward shift in myofiber size helps to account for higher muscle mass in PDGFRβ+. Because PDGF has been linked to insulin resistance (36), we wanted to determine whether overactive PDGFRα or PDGFRβ signaling altered insulin sensitivity or glucose handling. We showed no differences in GTT, ITT, or fasted blood glucose levels between PDGFR+ or CON, however, male PDGFRα+ had higher basal glucose levels compared to their CON (**Supplemental Figure 2A-H**).

Because PDGFRβ signaling can promote pericyte differentiation and angiogenesis (41) we investigated whether there was a change in capillary density in PDGFRα+ and PDGFRβ+ mice. Male PDGFRα+ mice had a higher capillary per fiber ratio compared to female PDGFRα+ mice (**Figure 2A-B**). However, the distribution of capillary number per myofiber was shifted left for male PDGFRα+ mice compared to CON, indicating a greater percentage of myofibers that had fewer capillaries compared to CON (**Figure 2C**). While PDGFRβ+ showed no differences in the capillaries per fiber or capillaries per area ratios, the capillaries per fiber distribution was shifted right, indicating a greater percentage of myofibers that had a higher number of capillaries compared to CON (**Figure 2D-F**). Due to muscle mass differences, we wanted to determine whether there were differences in muscle growth signaling and protein synthesis between genotypes. mTORC1 regulates muscle protein synthesis and growth. We examined mTORC1 signaling by probing for the mTORC1 downstream targets: 4EBP1, p70S6 Kinase, and rpS6 total and phosphorylated protein. We showed that total 4EBP1 protein was less in male PDGFRα+ mice compared to CON and that total and phosphorylated 4EBP1 protein was less in male PDGFRβ+ mice compared to CON (**Figure 3A-D**). However, there were no differences in p70S6 Kinase and rpS6 protein under any condition (**Figure 3A-D**). Due to minimal changes in downstream mTORC1 signaling, we next examined growth and differentiation factor 8 (i.e., myostatin). Myostatin is a negative regulator of muscle hypertrophy, and we found that myostatin plasma levels were lower in male and female PDGFRα+ and in male PDGFRβ+ compared to CON (**Figure 3E and F**). Last, to determine whether there was higher muscle protein synthesis we administered deuterium oxide (D_2_O) to the animals for the last 7 days prior to harvest. There were no differences in myofibrillar protein synthesis in any of the groups (**Figure 3G and H**).

**Figure 2.**
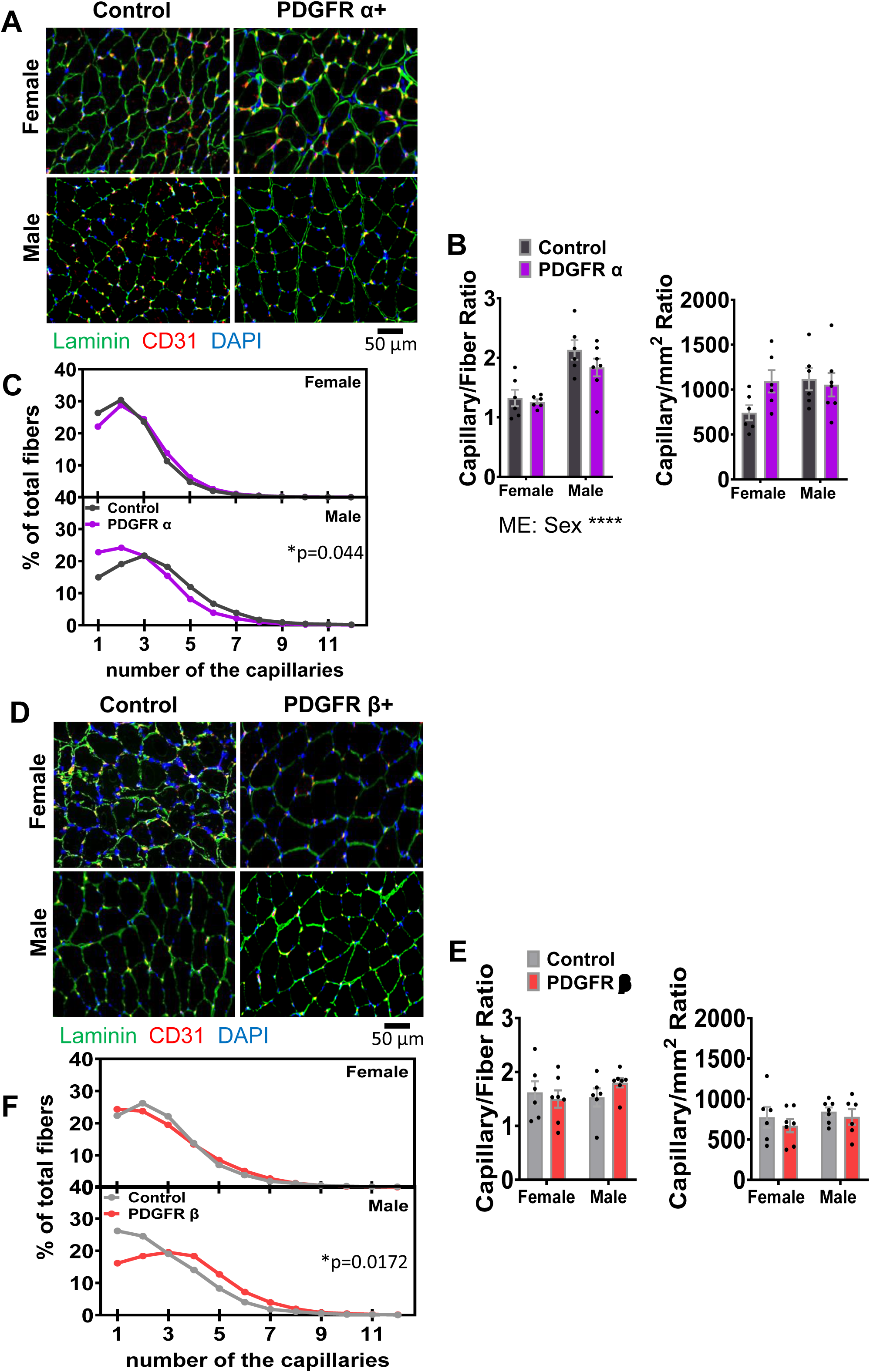
Overactive PDGFRα or PDGFRβ signaling only minimally effects capillary density. Representative images of GA muscle stained for capillarity (CD31) in **(A)** PDGFRα and **(B)** PDGFRβ mice. Associated distribution of capillary number per fiber in **(C)** PDGFRα and **(D)** PDGFRβ mice, and corresponding capillary/fiber and capillary/area ratios in **(E)** PDGFRα and **(F)** PDGFRβ mice (*n=6-7*). Data are means ± SEM. *p<0.05.

**Figure 3.**
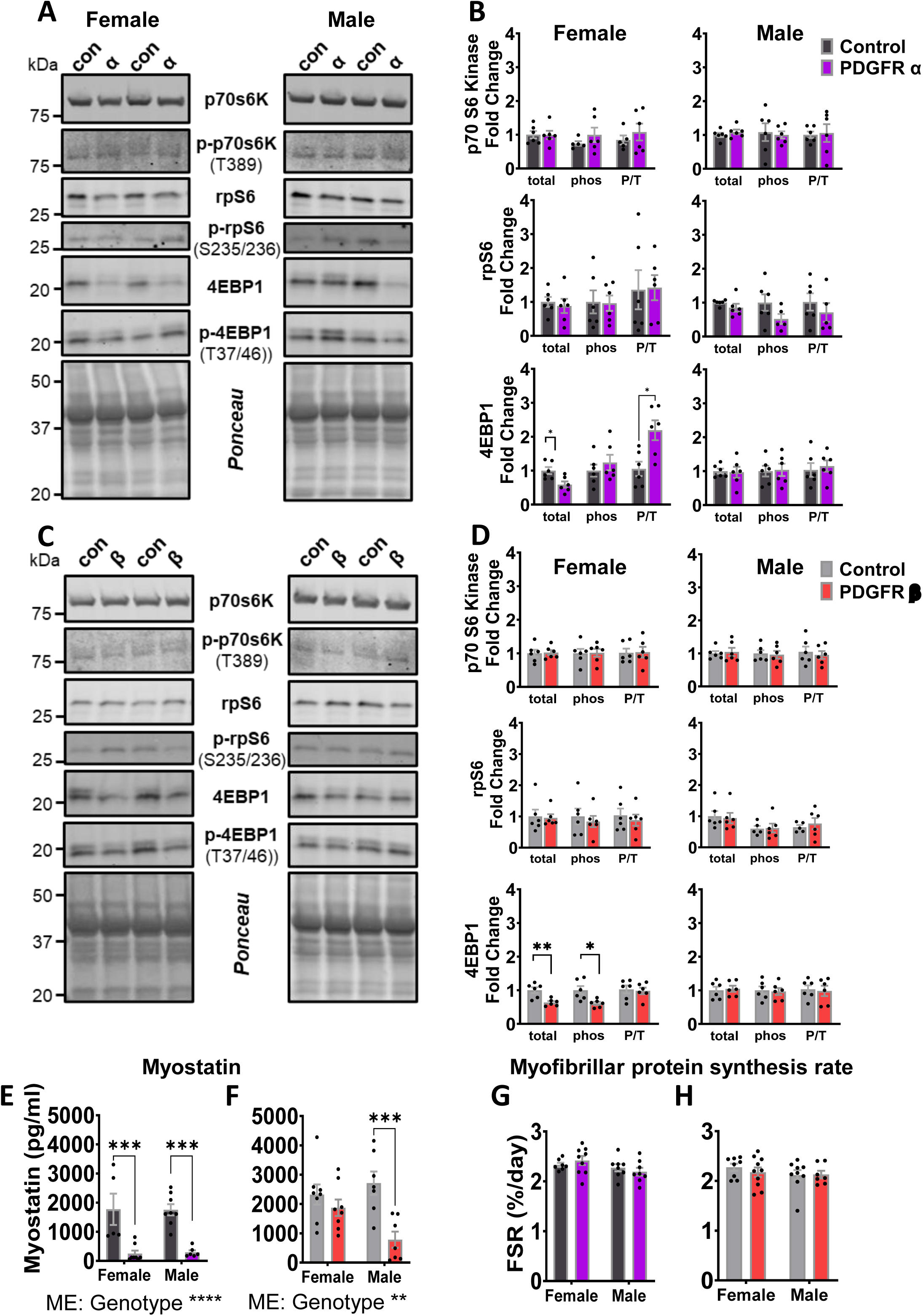
Mice with overactive PDGFRα or PDGFRβ signaling minimally effect mTOR signaling but have altered myostatin levels. Representative immunoblots and quantification of mTOR singling (phospho and total 4EBP1, p70s6K, and S6K) in TA muscle for **(A, B)** PDGFRα and **(C, D)** PDGFRβ mice (*n=7-8*). Plasma myostatin concentration for **(E)** PDGFRα and **(F)** PDGFRβ mice (*n*=6-9). Myofibrillar fractional synthetic rates (FSR) over 7-day labeling period prior to euthanasia in GA muscle of **(G)** PDGFRα and **(H)** PDGFRβ mice (*n*=7-9). Data are means ± SEM. *p<0.05, **p<0.01, ***p<0.001.

PDGFRα and PDGFRβ signaling potentially increases the synthesis of collagen from fibroblasts, FAPs, and skeletal muscle progenitor cells. To determine collagen abundance in skeletal muscle, we examined the amount of total, soluble, and insoluble collagen biochemically by measuring hydroxyproline content of these fractions in GA muscle. We first showed a main effect of sex and genotype for total, soluble, and insoluble collagen in PDGFRα mice. Second, we showed that male and female PDGFRα+ had more total and insoluble collagen compared to CON. Finally, we showed that female PDGFRα+ also had more soluble collagen compared to CON (**Figure 4A**). For PDGFRβ mice, there was a main effect of genotype for total and insoluble collagen abundance and a main effect of sex and genotype for soluble collagen abundance. Only total collagen was greater in female PDGFRβ+ compared to CON (**Figure 4B**). We also stained GA muscle cross sections with PSR. There was a main effect of genotype for densely packed collagen for PDGFRα+ mice, also female PDGFRα+ had a higher abundance of densely packed collagen compared to CON, while there was no difference in collagen packing for PDGFRβ+ compared to CON (**Figure 4C-F**).

**Figure 4.**
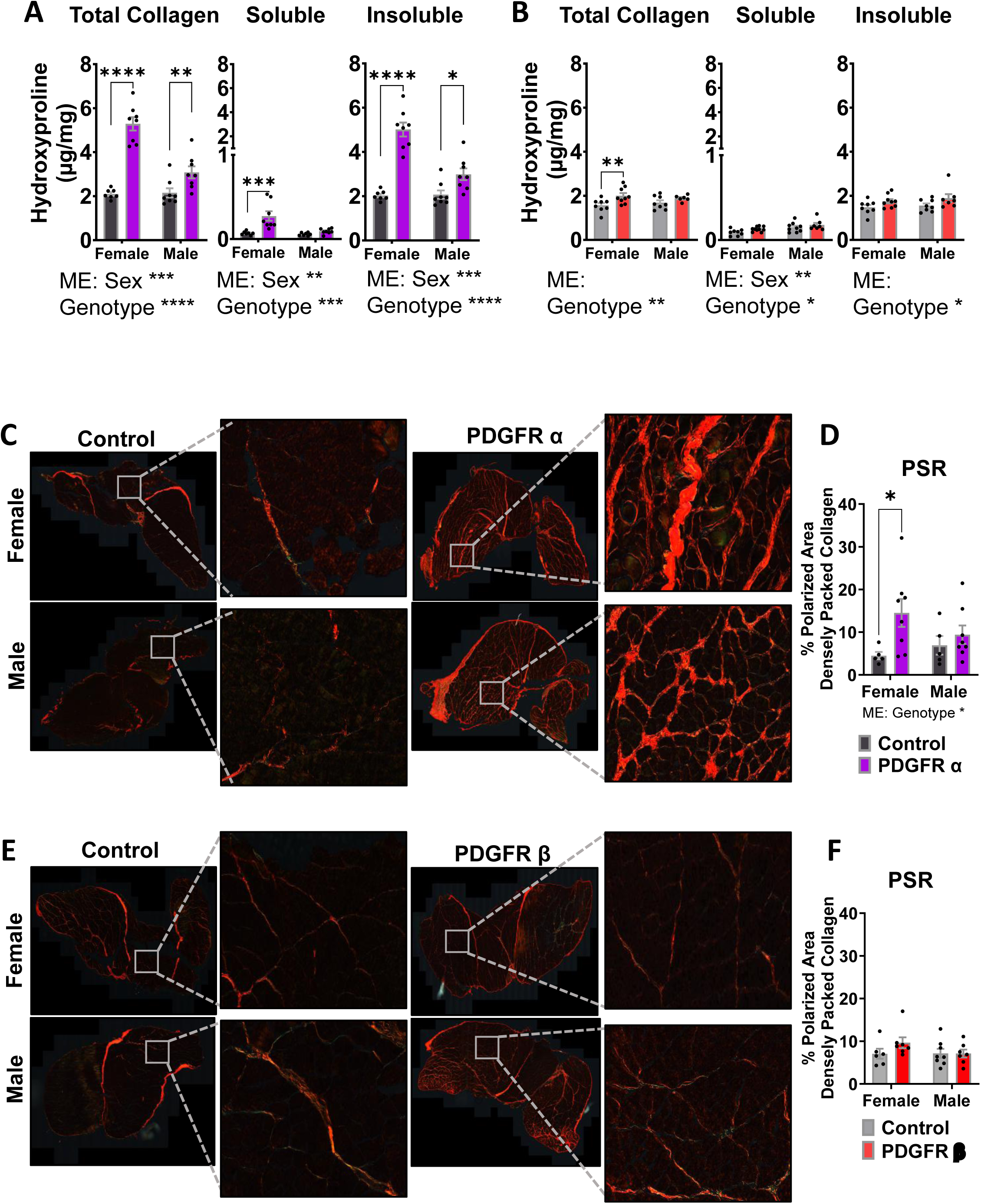
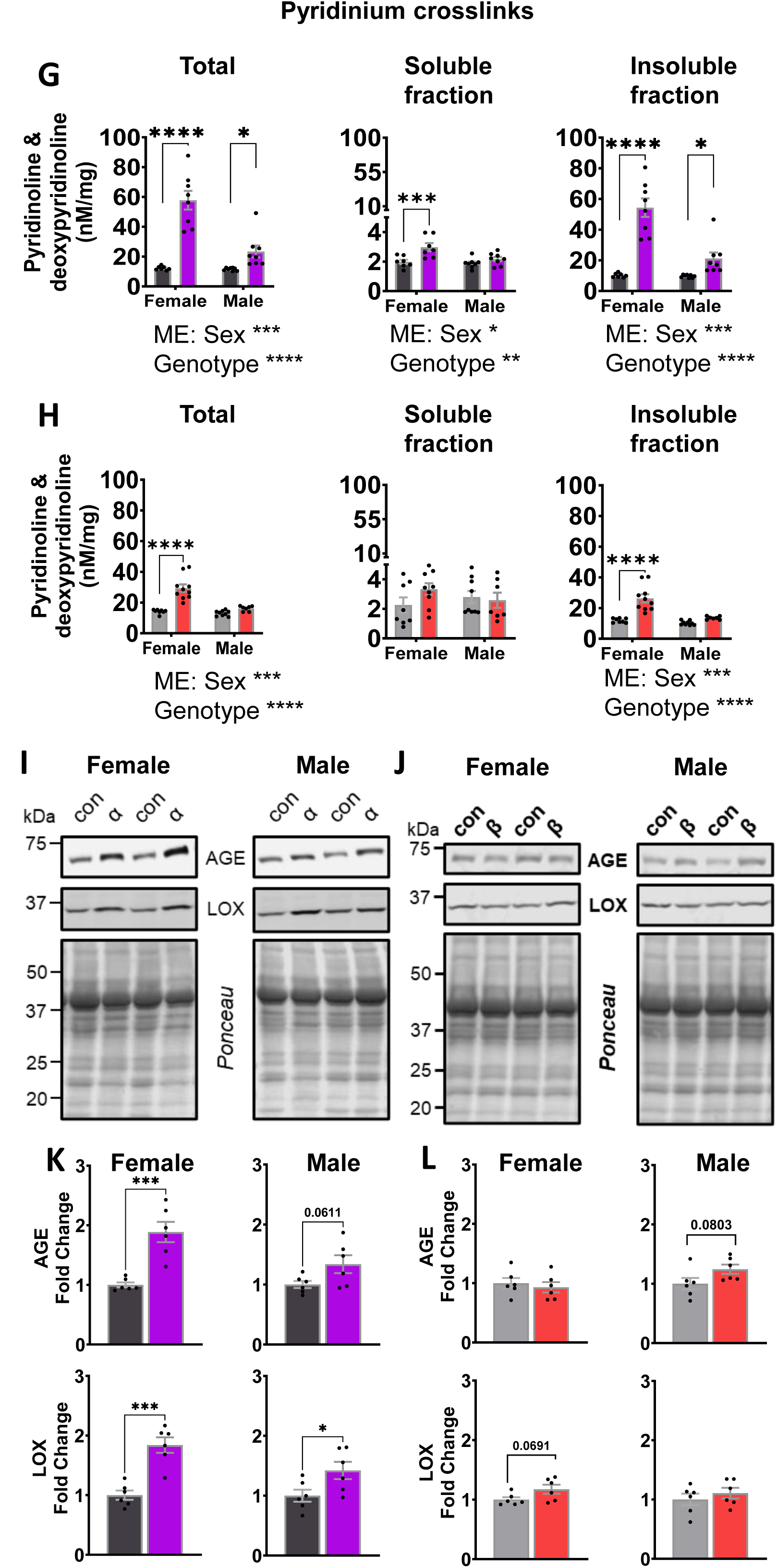
Overactive PDGFRα signaling results in a robust increase in insoluble and crosslinked collagen while overactive PDGFRβ model shows increase in collagen only in female mice. GA muscle was used for hydroxyproline, picrosirius red staining, and pyridinium crosslinks **(A-H)**. Total, soluble, and insoluble collagen concentration in muscle for **(A)** PDGFRα and **(B)** PDGFRβ mice (*n=7-8*). Representative picrosirius red (PSR) staining and collagen quantification in **(C, D)** PDGFRα and **(E, F)** PDGFRβ models (*n=7-8*). Total, soluble, and insoluble pyridinium crosslinks muscle for **(G)** PDGFRα and **(H)** PDGFRβ mice (*n=7-8*). Representative immunoblots and quantification of total AGE and LOX protein abundance in TA muscle for **(I, K)** PDGFRα and **(J, L)** PDGFRβ mice (*n=7-8*). Data are means ± SEM. *p<0.05, **p<0.01, ***p<0.001, ****p<0.0001.

Collagen crosslinking is a hallmark of fibrosis and decreases muscle function and force transmission. We showed main effects of P&D in total collagen and the insoluble collagen fraction of both PDGFRα and PDGFRβ groups. We also showed that P&D abundance in the total collagen fraction was greater in PDGFRα+ compared to CON and that this difference was entirely driven by higher abundance in the insoluble collagen fraction (**Figure 4G and H**). Female PDGFRβ+ had higher P&D in the total collagen fraction compared to CON, and although there was more P&D in the soluble collagen fraction of female PDGFRβ+, the majority of P&D was found in the insoluble collagen fraction (**Figure 4G and H**). As a final indicator of collagen structure/crosslinking, we measured LOX and AGE in TA muscles. We showed that both male and female PDGFRα+ had more LOX, and female PDGFRα+ had more AGE compared to CON (**Figure 4I and J**). There were no significant differences in LOX or AGE abundance in PDGFRβ+ compared to CON (**Figure 4K and L**).

We used several tests to determine how changes in fibrosis impact muscle function. First, we tested motor coordination using rotarod and saw no differences in either sex of PDGFRα+ compared to CON. There was, however, a main effect of genotype in which the genotype had shorter latency to fall (**Figure 5A**). PDGFRβ+ mice had a shorter latency to fall compared to CON in both sexes with a main effect of genotype (**Figure 5C**). We next performed maximal treadmill capacity and found that female PDGFRα+ ran a shorter distance before exhaustion compared to CON and had a main effect of genotype (**Figure 5B**). Similarly, female PDGFRβ+ mice also ran a shorter distance before exhaustion, and there were no differences in distance run with male mice (**Figure 5D**). There was a main effect of genotype in which the mutant genotype ran shorter distance before exhaustion (**Figure 5D**). It is important to note that each group ran to a similar level of exhaustion as indicated by the change in blood lactate levels with the treadmill test (**Supplemental Figure 3C and D**). Because body mass was different between mutant and CON mice, we measured max power output and total work performed, both of which incorporate body mass. We observed a main effect of sex for work performed (joules), as male mice had higher work output than female mice during the treadmill bout across both PDGFRα and PDGFRβ groups (**Supplemental Figure 3A and B**). There were no differences in maximal power output with PDGFR+ mice (**Figure 5B**). We showed that there was a main effect of sex and genotype regarding maximal power output in PDGFRβ+ mice, with males and PDGFRβ+ mice generating more power compared to female and CON respectively (**Figure 5D**). Female PDGFRβ+ mice generated higher maximal power compared to CON (**Figure 5D**).

**Figure 5.**
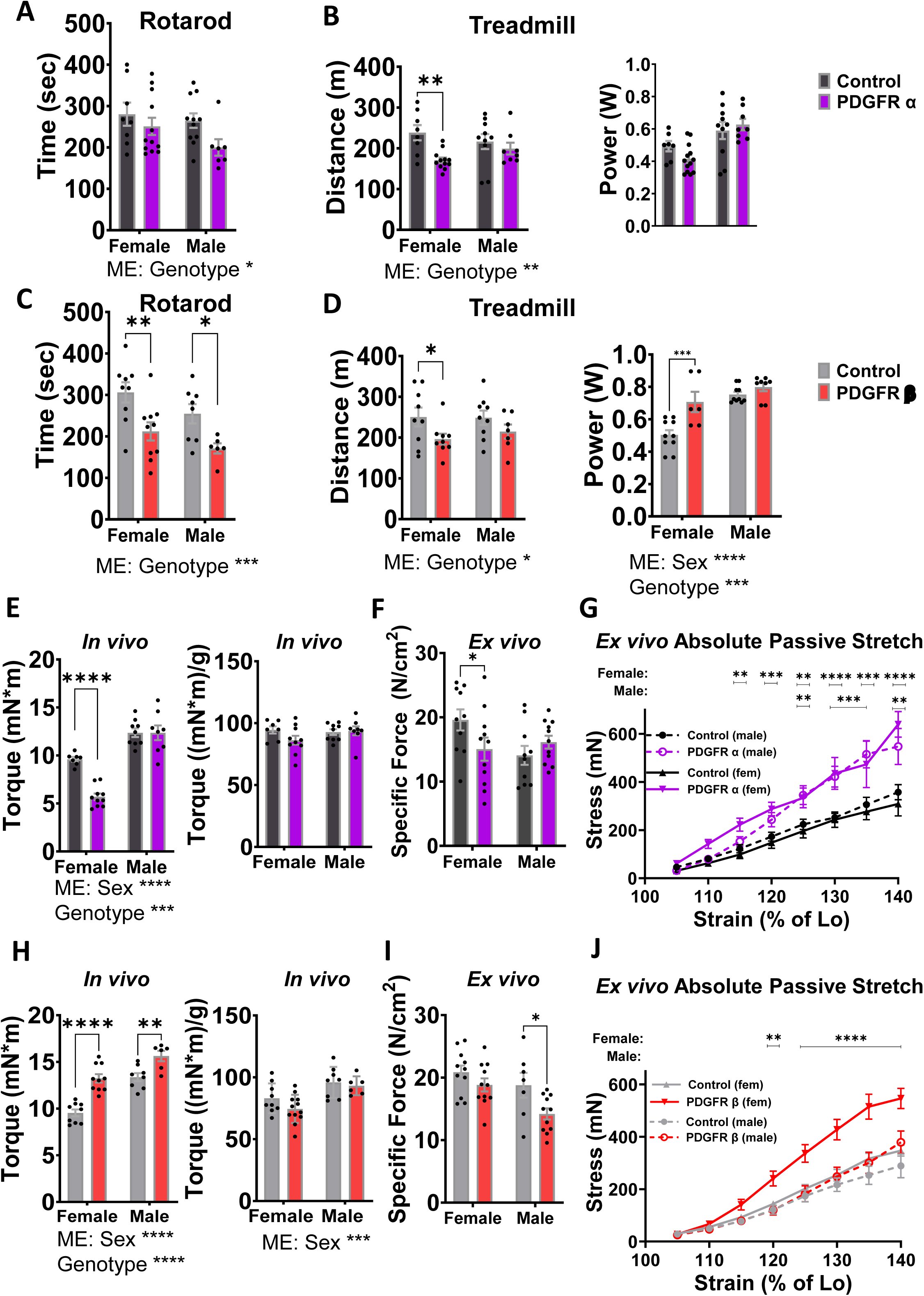
Overactive PDGFRα signaling results in stiffer muscles in both male and female mice while overactive PDGFRβ results in stiffer muscles only in female mice. Rotarod, treadmill run distance, and maximal treadmill power output for **(A, B)** PDGFRα and **(C,D)** PDGFRβ mice (*n=7-9*). *In vivo* maximal absolute and normalized (normalized to muscle mass) isometric torque for **(E)** PDGFRα and **(H)** PDGFRβ mice (*n=8-12). Ex vivo* maximal specific force and passive stretch values for **(F, G)** PDGFRα and **(I, J)** PDGFRβ mice (*n=8 12).* Data are means ± SEM. *p<0.05, **p<0.01, ***p<0.001, ****p<0.0001.

Last, we examined how overactive PDGFRα or PDGFRβ affected skeletal muscle strength and muscle stiffness. We first measured *in vivo* maximal force production of the plantarflexors (GA, PL, SOL) by electrically stimulating the tibial nerve. We showed that there was a main effect of sex and genotype in absolute *in vivo* torque and that values were lower in female PDGFRα+, with no differences observed in male PDGFRα+ compared to CON (**Figure 5E**). We also showed a main effect of sex and genotype for absolute torque, and that absolute torque was greater in both male and female PDGFRβ+ mice compared to CON (**Figure 5H**). When we normalized maximal *in vivo* torque to muscle mass, the main effect of sex remained but there were no differences between any PDGFRβ+ and their respective CON (**Figure 5H**). Further evaluation of the maximal torque curve showed that female PDGFRα+ had the slowest rate of contraction and relaxation, while both male and female PDGFRβ+ had higher rates of contraction and relaxation compared to CON (**Supplemental Figure 4E-F**). To determine the neural-independent specific force of the muscle, we used an isolated EDL in physiological solution and electrically stimulated the muscle to measure *ex vivo* contractile force. Female PDGFRα+ and male PDGFRβ+ had impaired force production compared to their respective CON (**Figure 5F and I**). To test muscle stiffness, we passively stretched the EDL muscle while simultaneously measuring force produced from the stretch. Male and female PDGFRα+ mice generated more force during passive stretch compared to CON in absolute (**Figure 5G**) and when normalized to muscle CSA (**Supplemental Figure 4C**), showing that PDGFRα+ have stiffer muscles. We also showed that female PDGFRβ+ generated more force with passive stretch compared to CON, however, we saw no differences in force generated during passive stretch test in male PDGFRβ+ (**Figure 5J**) or in either sex when normalized to muscle CSA (**Supplemental Figure 4D**). To gain greater insight into which aspects of collagen (i.e., total, soluble, insoluble, or crosslinked collagen) may contribute most to muscle stiffness, we correlated these outcomes with muscle stiffness. We show that for PDGFRα mice, insoluble, crosslinked insoluble, and total collagen had a relatively low, but significant positive correlation with muscle stiffness while soluble did not have a significant correlation (**Supplemental Figure 5A, B, D, F, H**). For the PDGFRβ mice, only crosslinked insoluble collagen had a low, but significant positive correlation with muscle stiffness (**Supplemental Figure 5A, C, E, G, I**).

## DISCUSSION

We hypothesized that both overactive PDGFRα or PDGFRβ signaling would increase skeletal muscle fibrosis through collagen deposition, leading to stiffer, and less functional muscle in mutant mice relative to their respective controls. We also hypothesized that overactive PDGFRα or PDGFRβ signaling would have sex-specific effects. Finally, we sought to determine if we either model would recapitulate age-related changes in fibrosis without using confounding interventions like muscle damage. Our results show that compared to CON, PDGFRα+ had a fibrotic phenotype as evident by more collagen deposition, increased collagen crosslinking, and higher AGE/LOX protein levels, which all correlated with greater muscle stiffness. Although PDGFRβ+ had minor changes in some markers of collagen deposition, there were notable changes seen were increased muscle mass, reduced fat mass, and prominent sex-specific differences in collagen crosslinking and muscle stiffness. In sum, these findings highlight the contrasting roles of PDGFRα and PDGFRβ in adult skeletal muscle, with PDGFRα acting as a potent inducer of collagen deposition and fibrosis, and a previously undescribed role of PDGFRβ in promoting skeletal muscle hypertrophy *in vivo*. These findings also identify PDGFRα+ as a mouse model to study skeletal muscle fibrosis in the absence of aging, muscle damage, or other disease process.

### Overactive PDGFRα Promotes Robust Fibrosis in Adult Skeletal Muscle

A notable finding of this study was the difference between the effects of overactive PDGFRα and PDGFRβ on collagen accumulation in skeletal muscle. Both male and female PDGFRα+ mice had a higher collagen concentration and amount of crosslinked collagen compared to CON. Overactive PDGFRβ resulted in only minor increases in collagen deposition and only in female mice. These results are important because an accumulation of more and stiffer/crosslinked collagen that is resistant to turnover is a hallmark of muscle fibrosis (7, 37).

With the increased fibrosis, there was a striking differences in muscle stiffness in male and female PDGFRα+ compared to their CON. To measure muscle stiffness, we passively stretched EDL muscles at increasing lengths and showed that male PDGFRα+ mice had a 53% increase in force while female PDGFRα+ mice had a 107% increase in force compared to their respective CON. Not only did these increases in insoluble and crosslinked collagen lead to stiffer muscles, but female PDGFRα+ mice had smaller type IIx and type IIb myofibers and lower specific force. However, we do not know if the changes in myofiber size are directly related to PDGFRα signaling, or secondary to fibrosis.

This study identifies PDGFRα over PDGFRβ activity as the primary driver of collagen deposition in adult skeletal muscle. Although we did not identify the mechanisms of how overactive PDGFRα promotes collagen deposition, previous studies have shown that PDGFRα+ cells differentiate into collagen type-I-producing cells that accumulate in fibrotic areas of muscle in *mdx* mice (35) and that constitutively active PDGFRα fibroblasts promote fibrosis during injury/repair (36). It is important to note that while previous studies demonstrate that PDGFRα promotes fibrosis in muscle during disease states or during injury/repair, there is a scarcity of data regarding the role of PDGFRα in skeletal muscle in the absence of damage or disease, and how PDGFRα activity may contribute to age-related fibrosis. Previous data from our lab indicate that the accumulation of collagen in muscle with denervation (37) and aging (27) are not due to higher rates of collagen synthesis, but rather a slowing of collagen breakdown. This slowing is likely caused by increased cross-linking that impairs breakdown (38, 39). From the current study, it is not clear if PDGFRα+ caused higher rates of collagen synthesis followed by increased cross-linking, or if PDGRα+ increased cross-linking that in turn slows breakdown and promotes fibrosis. When examining the concentration of crosslinks in proportion to collagen amount, there are higher proportions of crosslinks in PDGRα+ (57.8nM/mg in female and 23.4nM/mg in male) compared to CON (12.2nM/mg in female and 11.5nM/mg in male), particularly in female mice which had a ∼370% increase. The higher proportion of crosslinks to collagen concentration in PDGRα+ raises the possibility that PDGRα+ may also have a role in activating crosslinking. With either possibility, the overactive PDGFRα promotes fibrosis independent of disease or injury and thus could be used as a model to isolate the impact of fibrosis on skeletal muscle.

### Overactive PDGFRβ Leads to Skeletal Muscle Growth and Reduced Fat Mass

An unexpected finding was the effects that overactive PDGFRβ had on muscle mass and body composition. Compared to CON, PDGFRβ+ had higher overall body mass (38% higher in female and 20% higher in male mice) that was accompanied by substantial increases in lean mass (86% more in female and 49% more in male mice). In male, but not female PDGFRβ+ mice, this pronounced hypertrophy was associated with lower myostatin (GDF-8), a potent negative regulator of muscle growth (40). Interestingly, circulating myostatin levels were also reduced in both male and female PDGFRα+ mice, yet muscle mass was unchanged in males and lower in females compared to CON. This discrepancy demonstrates that low myostatin alone is not sufficient to induce muscle growth with overactive PDGFRα signaling, but the exact mechanism of how this occurs is unknown. PDGFRβ signaling can promote pericyte differentiation and angiogenesis (41), which was evident by a maintenance of capillarity in PDGFRβ+ with the greater muscle mass.

Overactive PDGFRβ mice had greater maximal power output than controls during treadmill running, possibly due to more muscle mass, but at the cost of impaired motor coordination. These findings of enhanced output but reduced coordination may reflect biomechanical challenges brought about by excess body mass (45) and altered limb mechanics, as indicated by the visibly widened stance and modified gait in PDGFRβ+ mice (data not shown). In addition to identifying different mechanistic actions through which these two receptors signal as highlighted above, these data identifiy distinct functions of PDGFRα and PDGFRβ in muscle function and stiffness.

PDGFRβ signaling is linked to both adipogenic and fibrotic fates. For example, in aged muscle, PDGFRβ lineage cells acquire mesenchymal stem cell-like properties and express both adipogenic and fibrotic markers (42). Given these diverse functions, activation of PDGFRβ signaling could influence adipogenic or fibrotic outcomes in PDGFRβ+ mice. We found that overactive PDGFRβ signaling led to a marked reduction in fat mass (53% in females and 48% in males) compared to CON. Another study found that adipogenesis is enhanced in PDGFRβ knockout mice (22). These data shed light on the regulatory role of PDGFRβ in maintenance of fat mass. These data identify a novel role for PDGFRβ in promoting skeletal muscle hypertrophy and reducing fat mass *in vivo*, potentially through myostatin signaling and/or other signaling pathways not investigated in this current study.

### Sex-Specific Differences Observed with Overactive PDGFRα and PDGFRβ

There were prominent sex-specific differences in this study for both PDGFRα and PDGFRβ groups. One significant sex difference was the large change in collagen composition. Female PDGFRα+ mice had higher levels of total, insoluble, and crosslinked collagen, as well as greater AGE and LOX content compared to male PDGFRα+ mice. Although less pronounced, similar trends were observed in female PDGFRβ+ mice, that had elevated P&D crosslinking compared to male counterparts. Female PDGFRβ+ mice also had stiffer muscles compared to male counterparts, likely due to higher P&D crosslinks. Despite higher P&D crosslinks in female PDGFRα+ mice, absolute muscle stiffness did not differ from male PDGFRα+ mice. However, when normalized to muscle CSA, female PDGFRα+ mice had higher stiffness compared to male PDGFRα+ mice. Another sex-specific finding was the lower muscle mass in the female PDGFRα+ compared to CON, whereas these differences were not noted in male PDGFRα+ compared to respective CON. Our data show that the smaller muscle mass in female PDGFRα+ was driven exclusively by lower type IIx and IIb myofiber size since type I and IIa fiber size was unaffected by PDGFRα+. Although direct regulation of sex hormones on PDGFR in skeletal muscle is lacking, our findings are consistent with the findings that estrogen and testosterone can modulate PDGF activity in other tissues and cell types (20, 46–48). Together, these findings underscore that there are sex-specific impacts of PDGFR-driven muscle remodeling.

### Limitations, Recommendations, and Conclusions

One limitation of the current study is that we measured protein synthesis and other biochemical assays (i.e., mTORC1 signaling, myostatin) at a point where rapid changes were likely completed and there was a new steady state in muscle mass and other collagen-related outcomes. We approached this first experiment as an endpoint analysis after a sufficient period of time for the overactive PDGFR activity to potentially have an effect. To capture rapid changes would require an additional cohort that were euthanized at an earlier timepoint. As a result, it is likely that our biochemical assays missed earlier changes that led to body composition changes in PDGFRβ mice (e.g., increased lean/muscle mass and decreased fat mass). Future studies could examine the time ∼2-6 weeks following tamoxifen induction to capture the mechanisms driving body mass changes as they occur. Another limitation is that our labeling approach did not allow us to capture muscle collagen turnover. We have shown that to properly assess collagen, especially when fibrosis is expected, there needs to be a time-course labeling approach (e.g., 1, 3, 7, and 14 days) (7, 27). This approach allows determination of valid synthesis rate because it does not assume that the collagen pool fully turns over, which we have repeatedly shown to be true (7, 27, 37). It is important to note that both PDGFRβ+ and PDGFRβ CON mice are Stat1 deficient, as overactive PDGFRβ signaling causes early lethality through Stat1-driven inflammation (25). Lastly, while our study focused on skeletal muscle, our mutant mouse models are not muscle specific, limiting our ability to attribute effects solely to muscle-resident cells. However, these models had outcomes that were consistent with receptor signaling in the muscle.

In conclusion, we showed that our hypothesis was partially supported in that overactive PDGFRα signaling led to fibrosis as evident by deposition of insoluble and crosslinked collagen and greater muscle stiffness. In contrast, overactive PDGFRβ had a sex-specific response with only female muscle expressing more crosslinked collagen and to a lesser degree than overactive PDGFRα. Surprisingly, overactive PDGFRβ increased muscle size and decreased fat mass. We saw sex-specific differences with fibrotic remodeling, muscle stiffness, and muscle size in responses to overactive PDGFRα and PDGFRβ signaling. PDGFRα+ mice showed multiple indicators of fibrosis, highlighting the value of the PDGFRα model for studying fibrosis in the absence of other confounding interventions (**Table 1**). Together, these findings establish PDGFRα and PDGFRβ signaling as distinct regulators of muscle remodeling and highlight the importance of considering sex-specific outcomes when investigating mechanisms of muscle fibrosis and aging.

**Table 1.** Summarizing the effects of overactive PDGFR α and β compared to aged muscle. Green boxes highlight differences between PDGFRα and PDGFRβ mice while purple boxes highlight sex differences.

| | | Overactive PDGF $\alpha$ receptor | | Overactive PDGF $\beta$ receptor | | Aged muscle |
| --- | --- | --- | --- | --- | --- | --- |
|  |  | ♀ | ♂ | ♀ | ♂ |  |
| Mass/ muscle mass |  | - | - | ↑↑ | ↑↑ | ↓ |
| Fat mass |  | - | - | ↓↓ | ↓↓ | ↓ |
| mTOR signaling |  | - | - | - | - | ↑ |
| Myostatin |  | ↓↓ | ↓↓ | - | ↓↓ | ↑ |
| Myofibrillar synthesis |  | - | - | - | - | ↑ |
| Collagen |  | ↑↑ | ↑↑ | - | - | ↑↑ |
| Pyridinium crosslinks |  | ↑↑ | ↑↑ | ↑↑ | - | ↑ |
| AGE/LOX |  | ↑ | ↑ | - | - | ↑ |
| Rotarod |  | - | - | ↓↓ | ↓↓ | ↓↓ |
| Treadmill distance |  | ↓ | - | ↓ | - | ↓↓ |
| Treadmill power |  | - | - | ↑ | ↑ | ↓↓ |
| Force |  | ↓ | - | ↑ | ↑ | ↓ |
| Force normalized |  | - | - | - | - | ↓ |
| Stretch normalized |  | ↑↑ | ↑↑ | - | - | ↓ |
| Stretch absolute |  | ↑↑ | ↑↑ | ↑↑ | - | ↓ |

## ACKNOWLEDGEMENTS

J.D.Fuqua was supported by NIA K99/R00 MOSAIC: 1K99AG086524-01A1 grant and the NIA Training Grant: 5T32AG052363-04. B.F. Miller a VA Research Career Scientist was supported by VA Merit Awards: VA I01 BX005592 and VA I01 BX006508. L.E. Olson was supported by R01-AR073828 and R01-AR080896. J.L. Brown was supported by a VA Career Development Award (1 IK2BX005620-01A1). Some images were created with BioRender.com. The authors declare no conflicts of interest.

## Supplemental Figures

**Supplemental Figure 1. Overactive PDGFRβ signaling results in greater muscle mass and longer tibia length** Body mass in **(A)** PDGFRα and **(B)** PDGFRβ mice at end of intervention period (*n=7-9).* Individual absolute wet muscle mass of the GA, PLA, SOL, TA, EDL muscles in **(C)** PDGFRα and **(D)** PDGFRβ mice (*n=7-9*). Tibia and EDL length for **(E, F)** PDGFRα and **(G,H)** PDGFRβ mice (*n=7-9*). Data are means ± SEM. *p<0.05, **p<0.01, ***p<0.001, ****p<0.0001.

**Supplemental Figure 2. Overactive PDGFRα and PDGFRβ signaling have no major impact on glucose handling and insulin sensitivity** Basal glucose levels for **(A)** PDGFRα and **(B)** PDGFRβ mice (*n=7-9)*. Fasted glucose levels, measured after 16h of fasting, for **(C)** PDGFRα and **(D)** PDGFRβ mice (*n=7-9).* GTT and ITT values for **(E,G)** PDGFRα and **(F,H)** PDGFRβ mice (*n=7-9)*. Data are means ± SEM.***p<0.001.

**Supplemental Figure 3. Mice from both PDGFR models produced similar work during treadmill exhaustion testing** Total work (joules) performed during treadmill tests for **(A)** PDGFRα and **(B)** PDGFRβ mice (*n=7-9).* Lactate levels, measured immediately after treadmill exercise, for **(C)** PDGFRα and **(D)** PDGFRβ mice (*n=7-9)*. Data are means ± SEM.

**Supplemental Figure 4. Overactive PDGFRα results in greater normalized passive stretch in female mice** Normalized force (normalized to CSA of muscle) generated by passive stretch for **(A)** PDGFRα and **(B)** PDGFRβ mice (*n=7-11).* Average rate of muscle contraction and relaxation for **(C)** PDGFRα and **(D)** PDGFRβ mice (*n=7-11).* Data are means ± SEM. *p<0.05, **p<0.01, ***p<0.001, ****p<0.0001.

**Supplemental Figure 5. Correlations between muscle stiffness and changes in muscle collagen content in mice with overactive PDGFR α and β signaling** Pearson r correlation heatmap between stiffness and variables of collagen content for **(A)** PDGFRα and PDGFRβ mice (*n=7-9).* Correlations between muscle stiffness and changes in total muscle collagen content in **(B)** PDGFRα and **(C)** PDGFRβ mice, soluble and insoluble collagen content in **(D,F)** PDGFRα and **(E,G)** PDGFRβ mice, and crosslinks in insoluble collagen fraction in **(H)** PDGFRα and **(I)** PDGFRβ mice (*n=7-9)*; Data are means ± SEM. *p<0.05.

## Notes

### Competing Interest Statement

The authors have declared no competing interest.

