## Supplemental Figures for "Overactive PDGFRα and PDGFRβ promote distinct yet overlapping phenotypes of skeletal muscle fibrosis and stiffness, with PDGFRβ also driving drastic muscle growth"

### Supplemental Figure 1

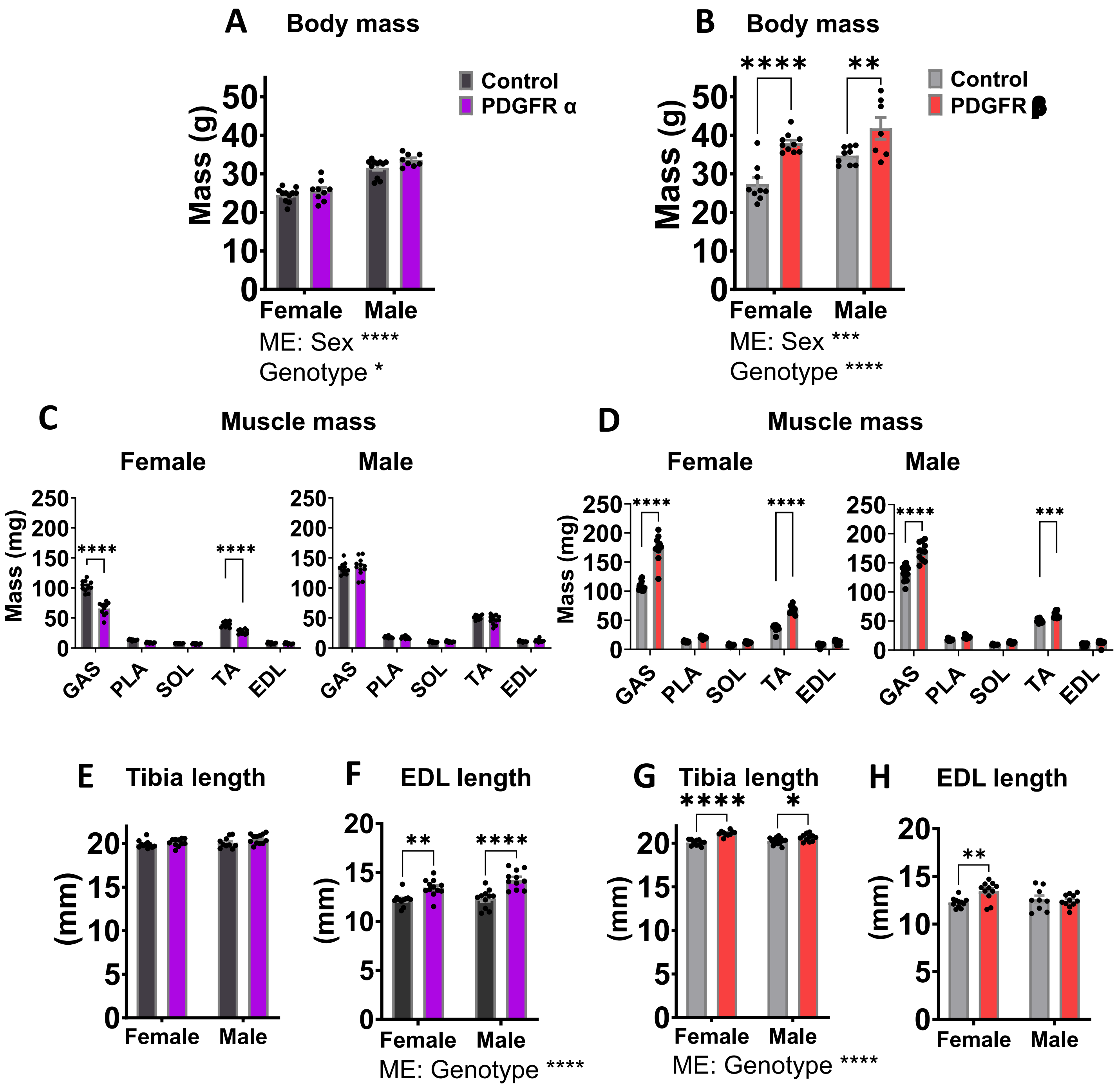

### Supplemental Figure 2

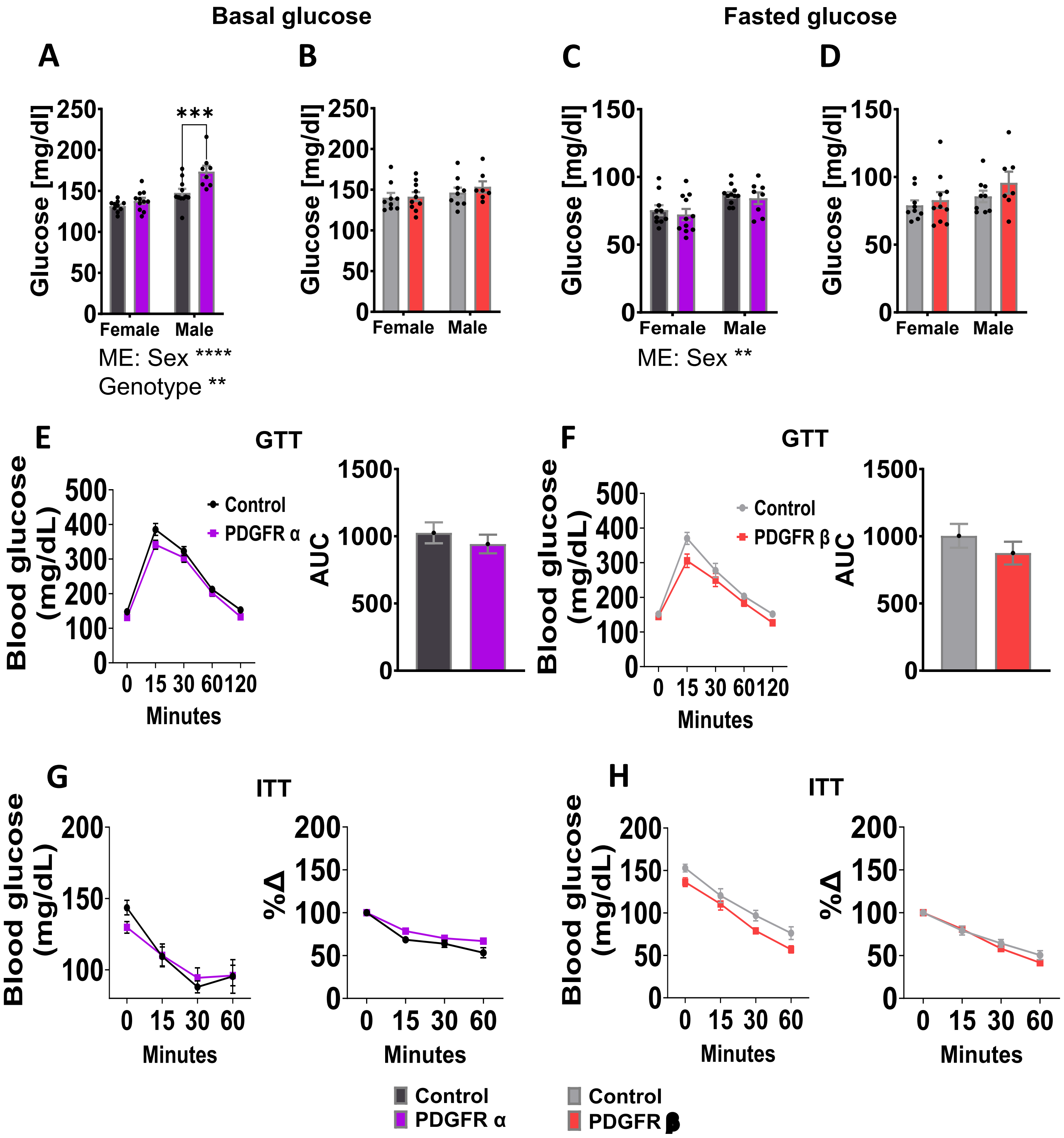

### Supplemental Figure 3

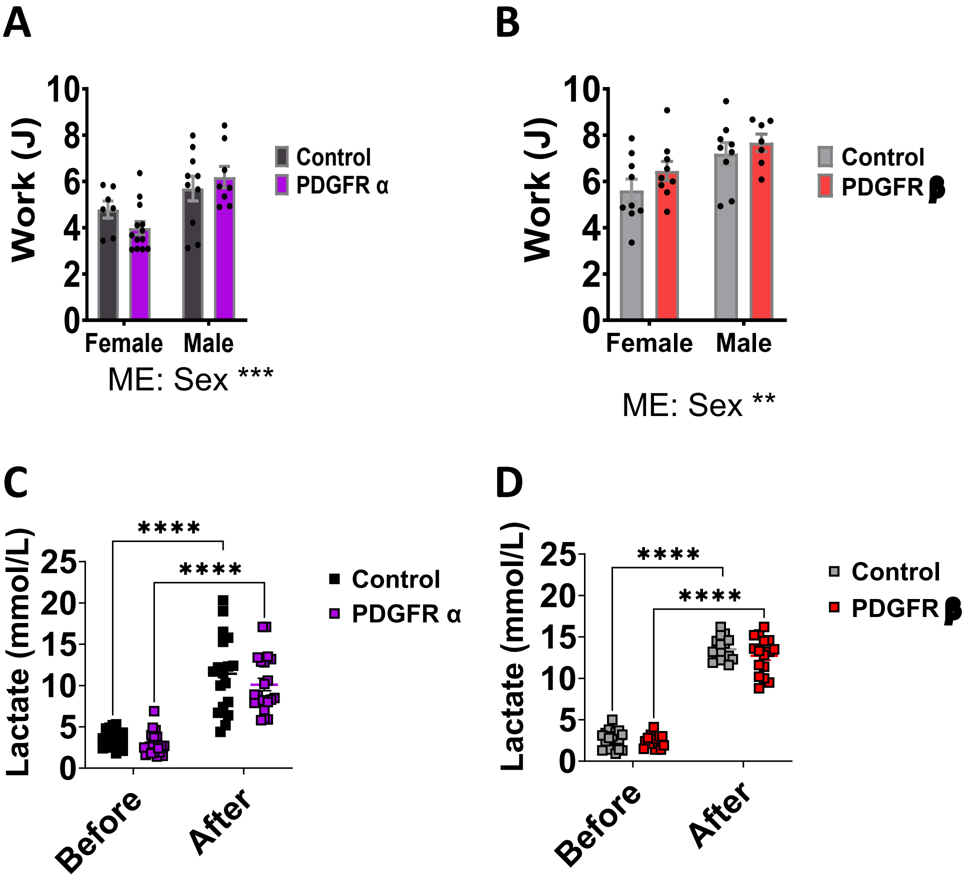

### Supplemental Figure 4

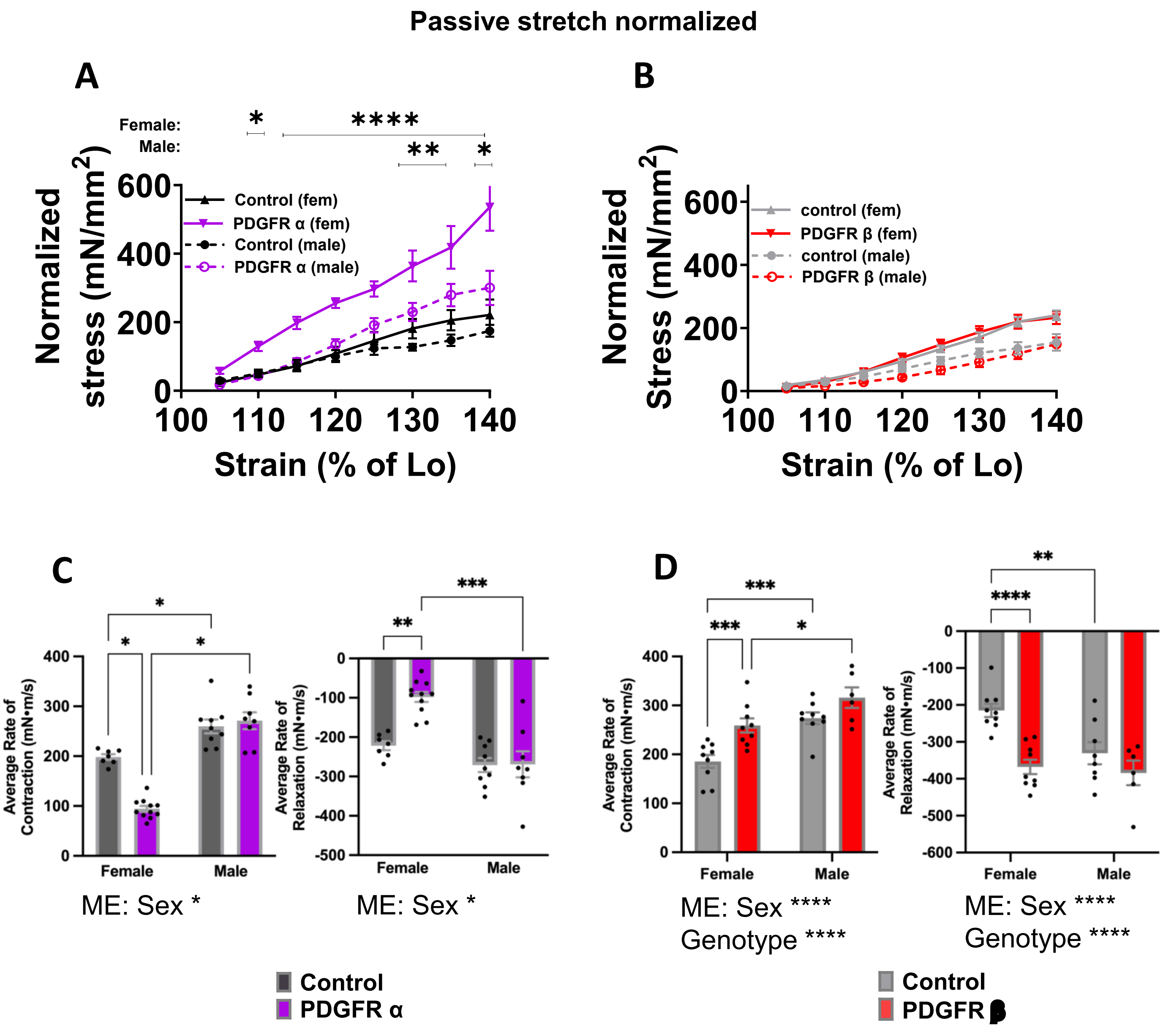

### Supplemental Figure 5

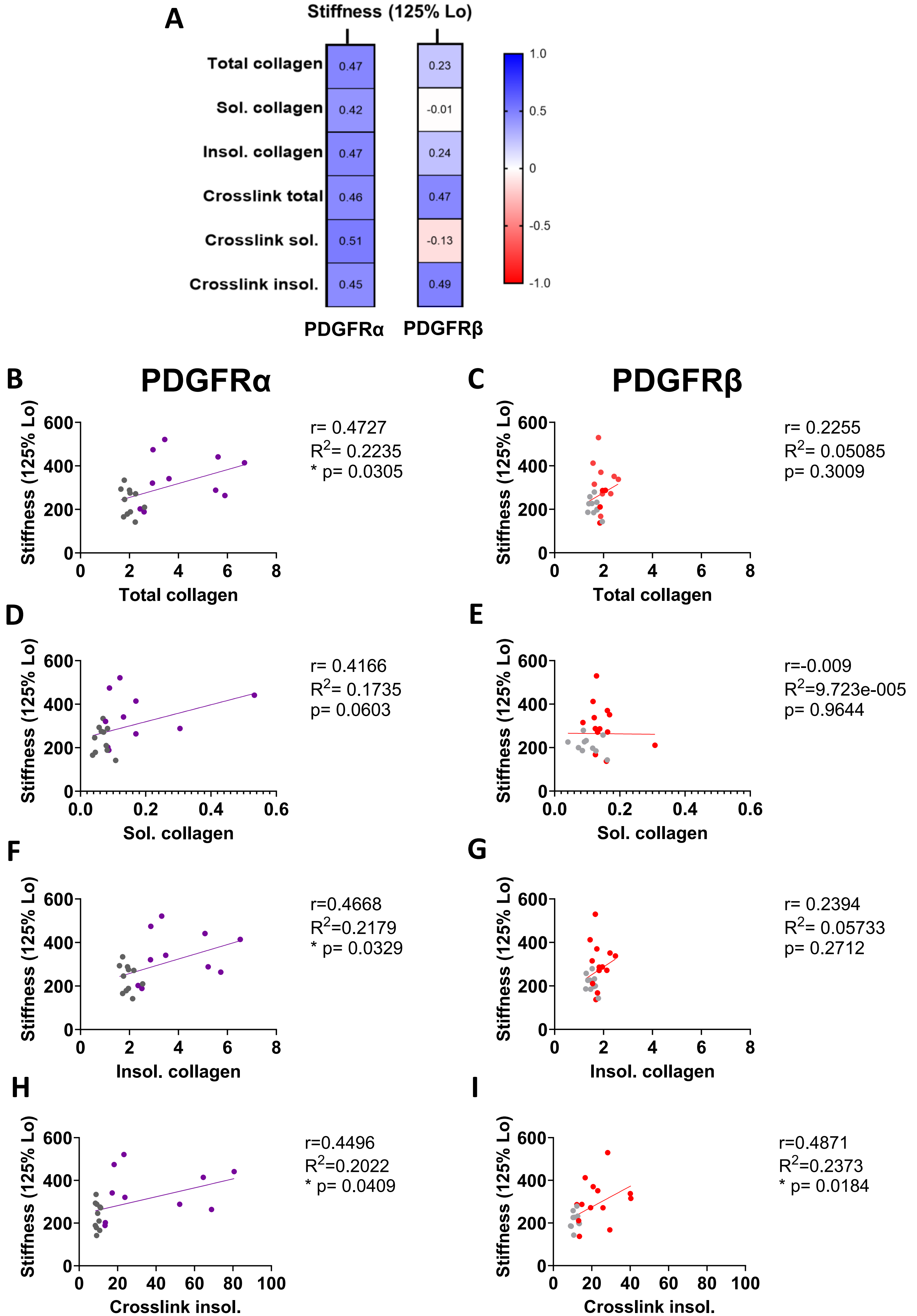
